# Loss of PR1 function enhances Arabidopsis resistance to *Botrytis cinerea*

**DOI:** 10.64898/2026.06.10.731468

**Authors:** Tamara Pečenková, Eva Kollárová, Tetiana Kalachova, Přemek Pejchar, Andrea Potocká, Anzhela Antonova, Radek Vitek, Tomáš Moravec, Lenka Burketová, Viktor Žárský, Martin Potocký

**Affiliations:** Institute of Experimental Botany of the Czech Academy of Sciences, Rozvojová 263, 165 02, Prague 6, Czech Republic; Department of Experimental Plant Biology, Faculty of Science, Charles University, Viničná 5, 128 44, Prague 2, Czech Republic

## Abstract

PATHOGENESIS-RELATED 1 (PR1) is one of the most widely used markers of salicylic acid (SA)-dependent plant immunity, yet its direct functional contribution to pathogen defence remains poorly understood. Here, we investigated the role of PR1 in *Arabidopsis thaliana* by analyzing a *pr1* loss-of-function mutant challenged with bacterial and fungal pathogens and fumonisin B1 (FB1)-induced cell death. Notably, loss of PR1 led to markedly different responses to distinct pathogens; while it moderately increased susceptibility to the pathogenic bacterium *Pseudomonas syringae,* it substantially enhanced resistance to the necrotrophic fungus *Botrytis cinerea*, and the responses to the necrotroph *Sclerotinia sclerotiorum* remained unaltered. The *pr1* mutant also displayed reduced spread of FB1-induced cell death, linking PR1 function to the promotion of stress-associated cell death. In line with the susceptibility changes, we observed the strongest PR1 accumulation and cell wall enrichment during *B. cinerea* infection using mCherry-tagged PR1 expressed under its endogenous promoter. Complementation with full-length PR1 and with a C-terminally truncated PR1 variant lacking the CAPE peptide restored wild-type susceptibility, whereas a non-cleavable PR1 variant did not. These results indicate that proteolytic processing at the CAPE cleavage motif, rather than the CAPE peptide itself, is required for PR1 function. Our data thus strongly suggest that PR1 may act as a susceptibility factor for necrotrophic pathogens by promoting host cell death.

## Introduction

Plant immunity relies on a multi-layered defence system comprising pattern-triggered immunity (PTI) and effector-triggered immunity (ETI), which activate overlapping responses including calcium influx, reactive oxygen species production, MAP kinase signalling, and induction of defence-related gene expression at both transcriptional and translational levels (Jones and Dangl, 2006; Ngou et al., 2021). These responses collectively function to counteract pathogen invasion, reinforce cellular barriers at infection sites, propagate systemic danger signals, and reprogram plant metabolic and developmental processes. A central hormonal regulator of these responses is salicylic acid (SA), which is primarily associated with resistance against biotrophic and hemibiotrophic pathogens and is a major inducer of systemic acquired resistance (SAR; Hayat and Ahmad, 2007; Raskin et al., 1992). This type of defence is closely linked to activation of a hypersensitive response (HR)-associated cell death, which is generally effective against biotrophic pathogens, but can be as well exploited by necrotrophic pathogens (Coll et al., 2011; Novaková et al., 2014). In contrast, jasmonate (JA)- and ethylene-dependent pathways are typically associated with defence against necrotrophic pathogens and act antagonistically to SA signalling (Ecker and Davis, 1987; Katsir et al., 2008).

PATHOGENESIS-RELATED 1 (PR1) is one of the most widely and historically used molecular markers of SA-dependent immunity and is strongly induced during pathogen infection and SAR activation (Jung and Hwang, 2000; Somssich et al., 1986; van Loon et al., 1999; Ward et al., 1991).

Although PR1 expression is an established proxy for immune activation, biological function of PR1 in plant defence remains incompletely understood. Members of the PR1/CAP (**C**ysteine-rich secretory proteins, **A**ntigen 5, and **P**R1 proteins; Gibbs et al., 2008) protein family have been implicated in antimicrobial activity and the modulation of immune responses. Evidence from diverse crop species suggests that PR1 proteins contribute to plant defence via immune signalling modulation (Breen et al., 2017; Carella et al., 2017; Chen et al., 2014, 2023; Kong et al., 2020;), and/or by the direct antimicrobial activity (Gamir et al., 2017; Guo et al., 2022; Han et al., 2023; Li et al., 2022; Luo et al., 2023; Pečenková et al., 2022; Riviere et al., 2008; Sarowar et al., 2005; Sung et al., 2021; Yang et al., 2018; Zhang et al., 2012, 2023; Zhou et al., 2021). It has also been found that the PR1 overexpression downregulates FB1-induced cell death (Lincoln et al., 2018). Notably, distinct mechanisms of PR1 action have been proposed across different plant–pathogen systems; however, none of these models fully accounts for all reported observations.

Given the limited number of studies addressing the biological function of PR1 in the model plant *Arabidopsis thaliana,* which are largely focused on subcellular localization and transient expression assays (Pečenková et al., 2017, 2022), we investigated the role of Arabidopsis PR1 by creation of the loss-of-function mutant *pr1*. Further on, by designing mCherry-tagged PR1 constructs we studied PR1 localisation in conditions of pathogen infections, and the functional significance of PR1-proteolytic processing. We demonstrate that loss of PR1 results in enhanced resistance to the necrotrophic fungus *Botrytis cinerea*, reduced sensitivity to fumonisin B1 (FB1)-induced cell death, and an increase in susceptibility to the bacterial pathogen *Pseudomonas syringae*. The infection by *B. cinerea* also led to the most pronounced increase in PR1 protein accumulation, further supporting a specific role for PR1 in defence responses against this pathogen. Collectively, these findings suggest that PR1 contributes to the regulation of host defence balance, potentially linking salicylic acid (SA)-associated signalling with the control of stress-induced cell death and pathogen-specific immune responses.

## Results and Discussion

### PR1 has a limited role in antibacterial defence but enhances susceptibility to *Botrytis cinerea*

To investigate the role of PR1 in response to patogens, we employed CRISPR/Cas9 technology and we created a new *pr1* loss-of-function mutat line (Supplementary table 1). The mutant has no obvious developmental or growth defect. We challenged *pr1* and WT plants with *Pseudomonas syringae* Pst DC3000 and we assessed disease development by colony-forming-units (CFU) quantification. To account for potential age-related differences in defence response, we used both flooding inoculation of plantlets and infiltration of adult leaves (Ishiga et al., 2012; Katagiri 2002). In both cases we found that the loss of PR1 function increases susceptibility compared with wild-type plants. More prominent difference was found for flooding of younger plants by ETI/HR-inducing avirulent AvrB strain, where the *pr1* mutant was found to be significantly more susceptible (Figure 1A). These results are in agreement with the proposed PR1 role in SA-related defence responses culminating with HR, however, the differences were only mild, suggesting that the PR1 function may be largely dispensable compared with upstream immune regulators.

**Figure 1.**
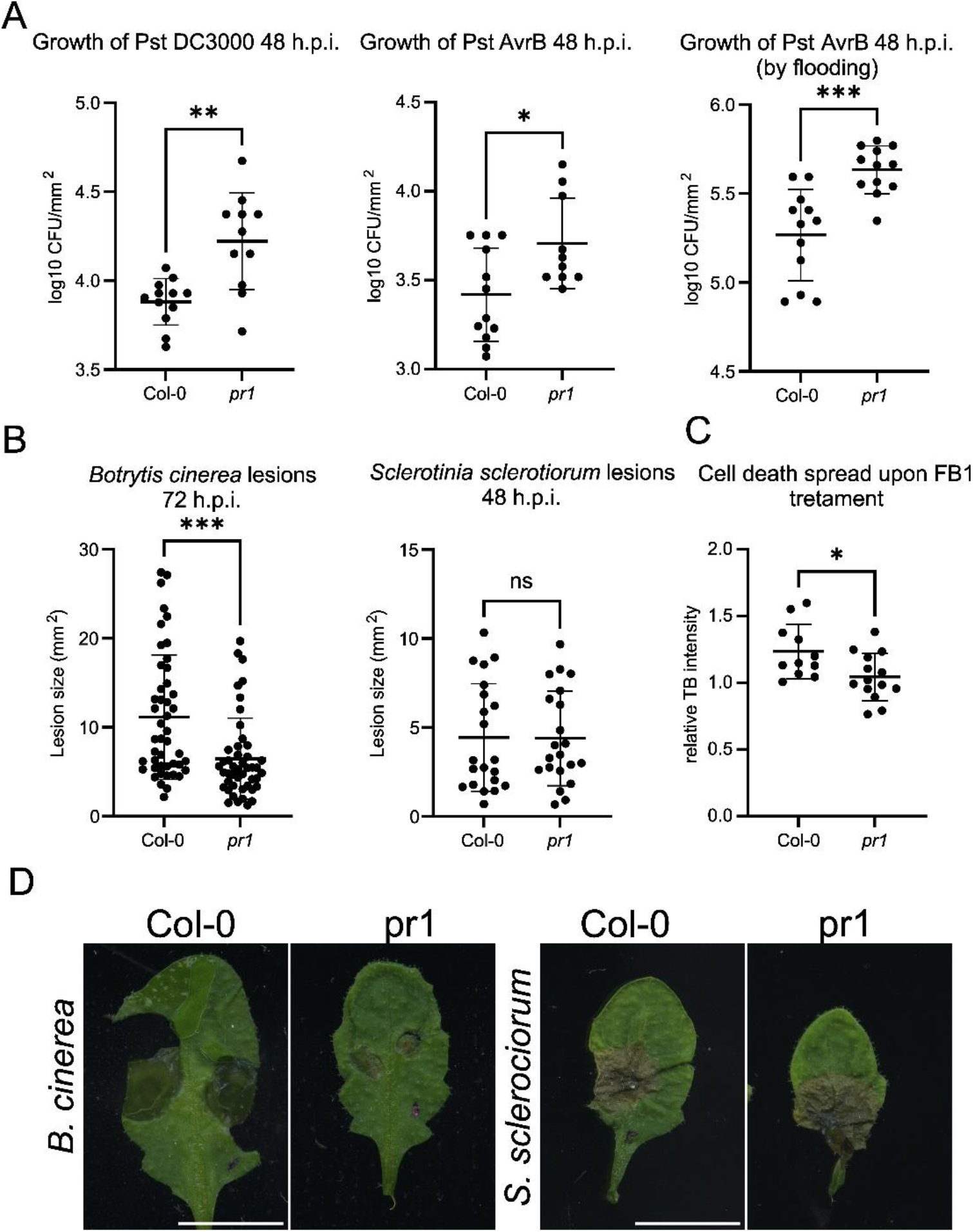
Loss of PR1 enhances susceptibility to pathogenic bacteria, but increases resistance to specific necrotrophic fungi. A) The *pr1* mutant is more sensitive toward bacteria Pst DC3000 and HR-inducing Pst AvrB. B) The sensitivity of *pr1* mutant toward *B. cinerea* is significantly decreased in comparison to WT (Col-0); resistance toward *S. sclerotiorum* is unaltered. C) The cell death spread upon FB1 infiltration of rosette leaves is more restricted in *pr1* mutant. (* - p<0.05, ** - 0.05<p>0.001, *** -p<0.001, determined by Welch’s t-test). D) Examples of B. *cinerea* and *S. sclerotiorum* lesions; bars, 1 cm.

In contrast to the relatively weak effect observed during bacterial infection of adult leaves, *pr1* mutant displayed substantially enhanced resistance to the necrotrophic fungus *Botrytis cinerea* (Figure 1B). Plants lacking PR1 displayed smaller lesions - less extensive disease symptoms, indicating that PR1 plays an important role in fungal colonization (Figure 1D). Notably, the response to another necrotrophic fungal pathogen, *Sclerotinia sclerotiorum*, was unaffected in the *pr1* mutant compared to wild-type (Figure 1B and 1D). Two non-mutually exclusive scenarios could explain this phenotype: PR1 may promote the spread of host cell death, which is required for efficient colonization by the necrotrophic pathogens, or it may act as a susceptibility factor specifically exploited by *B. cinerea* during infection. While the enhanced sensitivity of the *pr1* mutant to cell death-inducing AvrB bacterial strain is consistent with the first hypothesis, the unaltered responses to *S. sclerotiorum* suggest a more specific role for PR1 during *B. cinerea* infection.

To further test the role of PR1 in stress-associated cell death, we evaluated responses to FB1, a mycotoxin that induces defence-associated programmed cell death (Asai et al., 2000). PR1 accumulation frequently accompanies hypersensitive response-like cell death, although its functional contribution has remained uncertain (Betsuyaku et al., 2018). Loss of PR1 limited the spread of FB1-induced lesions, as visualized by Trypan-blue (TB) staining (Figure 1C; Supplementary figure 1). These findings indicate that PR1 positively contributes to the expansion of FB1-triggered tissue damage, and is in agreement with enhanced sensitivity toward the Pst AvrB and enhanced resistance toward *B. cinerea*.

Our findings are consistent with accumulating evidence that, although salicylic acid (SA)-dependent defences are traditionally associated with resistance to biotrophic and hemibiotrophic pathogens, components of the SA signalling pathway can also contribute to immunity against certain necrotrophic pathogens in a manner that depends on the stage of infection and tissue context (AbuQamar et al., 2006; Glazebrook, 2005; Hoppe et al., 2026; Vlot et al., 2009).

There are two possible explanations how PR1 could affect cell death spread - either PR1 actively participates in mechanical processes that facilitate lesion expansion, or the PR1 contributes to SA-defence responses activation until the death threshold is reached. In that case, missing PR1 function would prolong the time for tipping point. The PR1 dual role in pathogen defence and cell death regulation further highlights the complexity of this protein function during plant stress responses.

### PR1 expression and cell wall accumulation are induced by pathogens

By creating the mCherry tagged constructs placed under the control of endogenous promoter (Supplementary figure 2), we were able to analyse PR1 expression and subcellular localization during the pathogen-specific responses in *A. thaliana*. In rosette leaves of plants grown under the short day conditions, there was no observable signal of mCherry-PR1 expression unless biotic stress agents were applied. While *Pseudomonas* and *Sclerotinia* infection induced a modest and spatially limited increase in mCherry-PR1 signal (notably, mCherry-PR1 signal colocalized with the *Pseudomonas* colonies in the apoplast), *B. cinerea* triggered strong PR1 accumulation and a pronounced enrichment of PR1 in the cell wall compartment (Figure 2A). This localization pattern correlated with the stronger phenotypic contribution of PR1 during the interaction with *B. cinerea*.

**Figure 2.**
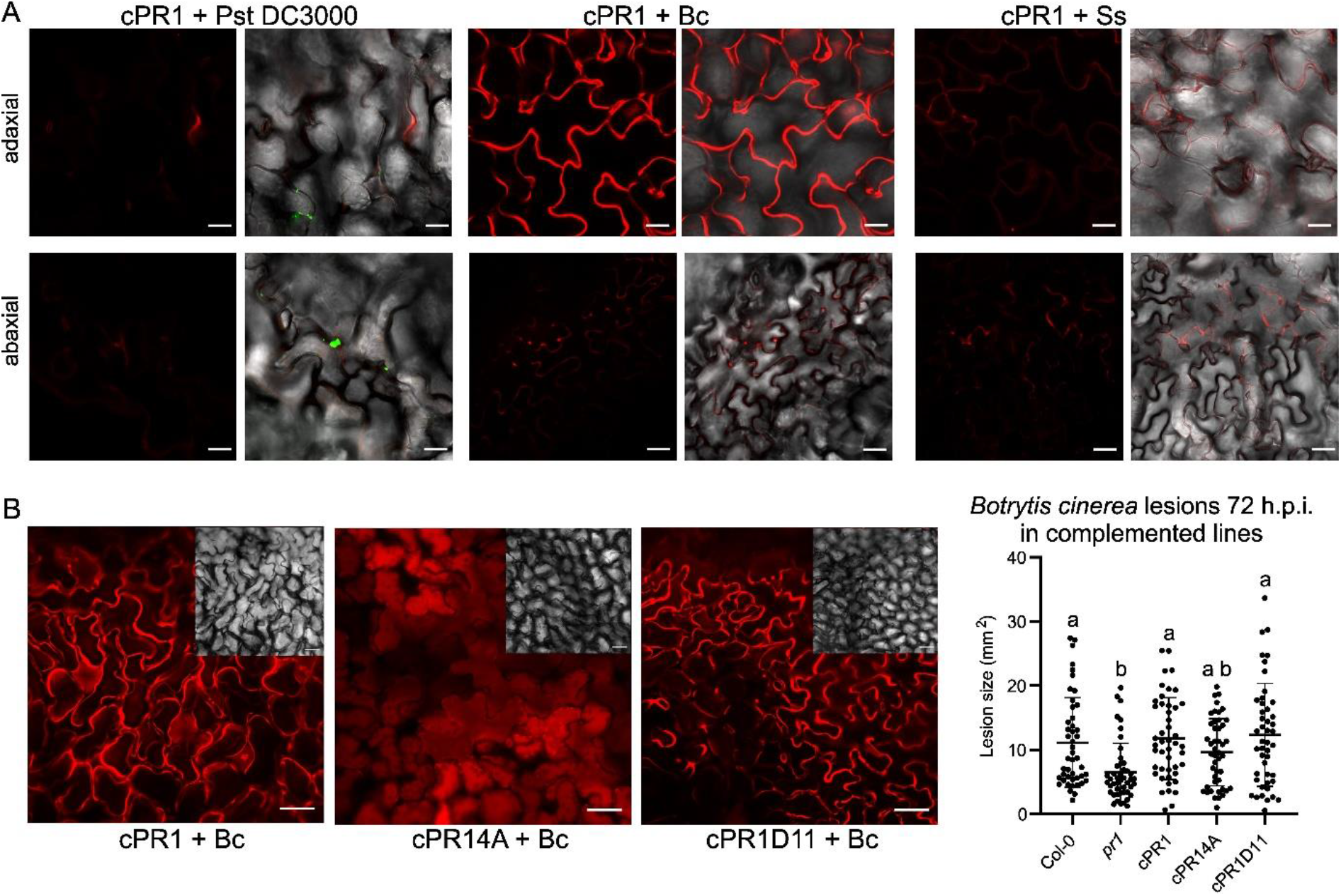
Localisation of PR1 during the pathogen infection and effects of mutant complementation on susceptibility toward B. cinerea. A) Accumulation of mCherry-PR1 in apoplast of leaves inoculated with GFP-carrying Pst DC3000, *B. cinerea* and *S. sclerotiorum*, both for abaxial and adaxial sides. Left panel – mCherry channel, right panels – overlay with GFP and bright field channels. Bars, 20 μm. B) Localization of *B. cinerea*-infected leaves, adaxial sides, in mutants complemented with full-length PR1 (cPR1), the CAPE cleavage site mutant cPR14A, as well as mutant variant devoid of CAPE peptide (cPR1D11). Bars, 50 μm. In the upper right corners are minimized appearances of the observed infected tissue on bright field. The chart shows the comparison of lesion sizes quantified for WT, mutant and complementation variants. Different letters indicate significant differences determined by ANOVA followed by Tukey post-hoc analysis, p<0.05, n=46.

These cell wall/apoplast observed localisations are consistent with a role for PR1 in the regulation or propagation of host cell death processes activated or exploited by *B. cinerea*. In general, pathogenesis-related (PR) proteins are secreted into the apoplast, where they are strategically positioned to interact with invading pathogens, particularly during the early stages of infection (rev. in Dos Santos and Franco, 2023). The prominent accumulation of PR1 in the cell wall during *B. cinerea* infection is in agreement with gene expression study by Jeblick et al., 2022, and points to its strong involvement in modification of tissues undergoing pathogen invasion and exposure to fungal cell wall-degrading enzymes (Espino et al., 2010; Voxeur et al., 2019). It is possible that distance fractions of PR1 protein produced in infection context contribute to local and systemic immunity. The potential systemic production of PR1 was unfortunately not possible to visualize with our microscopy-based experimental approach. It remains to be further explored whether the role of PR1 in systemic acquired resistance (SAR)-related signalling also contributes to sensitivity toward B. cinerea. (Breen et al., 2017; van Loon et al., 2006).

### Proteolytic processing and coupled secretion of PR1 contribute to sensitivity to *B. cinerea*

To determine whether proteolytic processing of PR1 releasing C-terminal CAPE peptide (Chen et al., 2014, 2023; Pečenková et al., 2022; Zhang et al., 2025) contributes to outcome of fungal infection, we employed in complementation of *pr1* mutant, besides the full-length PR1 (cPR1) also the variants carrying mutations that prevent cleavage of the C-terminal CAPE peptide (cPR14A), and a mutant variant devoid of CAPE peptide (cPR1D11; Pečenková et al., 2022; Figure 2B, Supplementary figure 2). In contrast to cPR1 and cPR1D11, the non-cleavable PR1 construct, despite being strongly induced by *B. cinerea* infection, did not localize to the apoplast but accumulated in the vacuole and failed to fully restore wild type response to *B. cinerea*, while two other variant complemented the mutation (Figure 2B and Supplementary figure 3). These results indicate that PR1-mediated resistance requires intact cleavage or cleavage mimicking for successful secretion, and not the intrinsic function of released CAPE peptide (Chen et al., 2023; Pečenková et al., 2022). By placing the fluorescent tag on the N-end of protein (behind signal peptide for entry into the secretion pathway), we prevented mixing of signals coming from the full-length protein and cleaved-off CAPE peptide, as it would be the case in the C-terminal fusion with fluorescent tag, meaning that the localisation and dynamics of CAPE peptide in this experimental setup remain obscure, and is something that we would like to address in future.

The strong requirement for CAPE processing and efficient secretion of PR1 during *B. cinerea* infection is consistent with our observation that PR1 accumulation and cell wall localization are the most pronounced in response to this pathogen. A local generation of CAPE peptides at infection sites may enhance defence signalling and contribute to cell death threshold. The possible contribution of CAPE peptide to the outcome of *B. cinerea* infection thus remains to be elucidated.

## Conclusion

The contrasting responses to different pathogens observed in *pr1* mutant suggest that PR1 protein contributes to the balance between distinct defence outputs rather than acting as a canonical positive regulator of immunity. A set of complementary experiments indicate that PR1 positively contributes to stress-induced cell death spread, such as increased resistance of *pr1* mutant to necrotrophic fungus *B. cinerea,* enhanced sensitivity to *Pst*, and reduced susceptibility to FB1-induced cell death. Our observations, however, conflicts report on cell death rescue by PR1-overexpression by Lincoln et al., 2018, which may be explained by the developmental stages-specific effects of PR1 (Pečenková et al., 2022). The PR1 thus contributes to the situation from which necrotrophic pathogens benefit. In contrast, the increase in susceptibility to *Pst* is consistent with a moderate reduction in SA-associated defence output, which is critical for restricting bacterial growth. Together, these observations indicate that PR1 influences defence signalling and the regulation of stress-induced cell death, with pathogen outcomes depending on whether host cell death promotes or restricts infection.

## Material and Methods

### Plant cultivation

For cultivation of plants, seeds were surface sterilised (5 min in 70% ethanol, 2 × 5 min in 10% commercial bleach, rinsed three times in sterile distilled water) and stratified for 2–3 days at 4°C. Seeds were then germinated and grown in Jiffy Products International pellets for 5 weeks at 21 °C and 10/14 h of light per day in growth rooms. As a wild type control, the Columbia-0 ecotype was used.

### Generation of CRISPR mutant line

The *pr1* mutant was created using CRISPR technology (Lei et al., 2014; Xing et al., 2014). An expression vector harbouring a two-target gene-specific gRNA has been constructed using reverse and forward primers for two PR1 gene-specific gRNAs (Supplementaary table 1). Target sequence fragments were amplified from pCBC-DT1T2 using Q5 polymerase (New England Biolabs) according to the manufacturer’s instructions and cloned into the vector pHSE401E using the Golden Gate cloning reaction with enzymes from New England Biolabs. The resulting construct expressing two gRNAs was used to transform *A. thaliana* Col-0 by standard *Agrobacterium tumefaciens* infiltration (Clough and Bent, 1998). Transgenic plants were selected by hygromycin resistance and presence of homozygous single base insertion was verified by sequencing. Homozygotes lacking the Cas9 fragment identified by PCR were used for further experimenting.

### Cloning of constructs

All PCRs were performed using Q5 polymerase (New England Biolabs). To create construct for pPR1::SP-mCherry-PR1 (cPR1), first entry clone was prepared as follows: the AtPR1 promoter (2342 bp upstream of startcodon) together with PR1 signal peptide sequence was amplified from Arabidopsis Col-0 genomic DNA using specific primers flanked by EcoRI and NcoI sites. Next, mCherry with eliminated middle NcoI site was commercially synthetised (Eurofins Genomics) flanked on the start of mCherry by NcoI site. Sequence of mature PR1 was amplified from Arabidopsis Col-0 genomic DNA using specific primers flanked by overhang to mCherry (containing BsrGI site) and XhoI site. Finally, all parts were introduced into pENTR3C (Invitrogen) vector by four-way ligation via EcoRI and XhoI sites. To prepare entry clones of mutated variants pPR1::SP-mCherry-PR14A (cPR14A), and pPR1::SP-mCherry-PR1D11 (cPR1D11), megaprimer strategy was used. First, reverse megaprimers containing mutated parts of PR1 region flanked by XhoI site were amplified using specific primers from pAtPR1::SP-mCherry-PR1 entry clone as a template. Next, megaprimers were used together with forward primer flanked by overhang to mCherry to amplify mutated mature PR1 regions using previously described entry clones PR1-4A and PR1-D11 as the templates (Pečenková *et al*., 2022). These regions were used to replace the original mature PR1 sequence in pAtPR1::SP-mCherry-PR1 via BsrGI and XhoI sites by three-way ligation (together with BglII site in PR1 promoter) to produce entry clones of mutated PR1. All entry clones were recombined by LR Clonase II (Invitrogen) into the Gateway destination vector pGWB601 (Nakamura et al., 2010) to generate final vectors. Created constructs were used for transformation of *pr1* mutant plants using floral-dip method (Clough and Bent, 1998).

### Pathogen assays

*Botrytis cinerea* B05-10 (Zimmerli et al., 2001) spores were kept in -20°C freezer at 10^6^ spores/mL stock in dH_2_O. Stock of spores was first diluted in dH_2_O (1:9, v:v) and then diluted 1:1, v:v in 50% PDB medium, to reach final concentration of 5×10^4^ spores/mL in 25% PDB. For the control 25% PDB was used. Two 5 µL drops of spore suspension or control were placed at adaxial side of fully mature leaves. Plants were kept at 100% humidity for 72 h (3 dpi) and then scanned (Epson Perfection V700 Photo, Suwa, Japan, at 1200 dpi resolution). Lesion area was measured by FiJi software (Schindelin *et al.*, 2012). At least three individual plants per treatment were used, three leaves per plant.

*Sclerotinia sclerotiorum* isolate Ss 05/08 (Ss 05) (Novakova et al, 2014) were cultivated at SNA medium for 4 days in the dark at room temperarture. Leaves were inoculated by placing one agar block (3 mm diameter) with young mycelium on the leaf surface into the 5 µL drop of dH_2_O. Plants were kept overnight at 100% humidity, and then scanned (Epson Perfection V700 Photo, Suwa, Japan, at 600 dpi resolution). Lesion area was measured by FiJi software (Schindelin *et al.*, 2012). At least three individual plants per treatment were used, three leaves per plant.

*Pseudomonas syringae pv. tomato* (Pst) DC3000 *–* Flooding assay was performed with modifications according to Ishiga et al. 2011. Briefly, we used *P. syringae pv. tomato* DC3000, and avirulent strain Pst AvrB (kindly provided by Michael Wrzaczek (Nimchuk et al., 2000)), OD = 0.01, diluted in water with Silwet (0.0025%), for flooding of 2-week-old plants for 2 min. Suspension was decanted from plates and seedlings incubated for 24 h on 12/12 h (day/night). After the incubation, plants were weighted, surface sterilized by 70% ethanol and homogenized by beads homogenizer. Series of dilutions were plated onto LB with rifampicin (25 µg/ml) and kanamycin for AvrB (25 µg/ml) and left for approximately 30 h until the colonies become visible and countable. The number of colonies with accompanying statistics was analysed and graphs created in GraphPad program.

The inoculation of *A. thaliana* by Pst and AvrB was performed according to previously adapted protocol (Kalachova *et al.*, 2023) : bacteria were cultivated overnight on plates containing LB medium (tryptone 10 g/L, NaCl 10 g/L, yeast extract 5 g/L, pH=7.0) supplemented with 1.4% agar and 50 mg/L rifampicin. Four-week-old plants (three leaves at the similar developmental stage, middle age leaves: 8th-9th-10th leaves) were syringe-infiltrated with the suspension of Pst (OD_600_=0.0005 in 10 mM MgCl_2_). At 3 days post inoculation, 3 discs (6 mm in diameter) were sampled from inoculated leaves from each plant, pooled (one plant as one sample) and homogenized in 1 mL of 10 mM MgCl_2_, in a 2 mL Eppendorf tube, with 1 g of 1.3 mm silica beads using a FastPrep-24 instrument (MP Biomedicals, USA). The resulting homogenate was subjected to serial 10× dilutions and pipetted onto LB plates. Colonies were counted after 2 days of incubation at 26 °C. Six individual plants were used per treatment; bacterial load being expressed as log10(CFU*mm^-2^) of leaf tissue.

### Preparation of GFP-expressing Pst

Plasmid for constitutive expression of GFP in bacteria was assembled using GoldenBraid cloning (Sarrion-Perdigones et al., 2013). The low copy number plasmid pLX-B2 alpha1 with a broad host range Ori for bacterial replication pBBR1 was chosen as backbone (Pasin *et al.* 2017). As the regulatory sequence, the bacterial promoter JM23100 from previously described Anderson’s promoters (Bhat *et al.* 2024) and HisOperon terminator (Di Nocera *et al.* 1978) were used. The AvGFP from the genebank collection of Cloning Facility IEB CAS was used as fluorescent marker. Previously described cloning protocol (Sarrion-Perdigones *et al.* 2011) was used with following modifications. In 15 μl reaction, 1x ligase buffer, 5U T4 DNA ligase, 5U BsaI, 0,2 μg/mL nuclease-free BSA, 60 ng of empty level-1 plasmid, 1 ul of each DNA part (each concentration about 80-100 ng/uL). After 15 min of initial restriction at 37°C, start 30 steps of cycling 37°C for 2,5 min and 16°C for 4 min. Final restriction took 1h at 37°C and inactivation 15 min at 80°C. The cloning reaction was transformed into chemically competent *E. coli* TOP10 cells by the heatshock protocol (Singh *et al.* 2010). Transformants were selected on LB plates containing kanamycin (50 μg/mL). Plasmid DNA was isolated using the QIAprep Spin Miniprep Kit and was verified by restriction with EcoRI and sequencing (Eurofins Genomics). Restriction enzymes were purchased from New England Biolabs, and T4 DNA ligase from Thermo Scientific.

Electrocompetent *Pst* cells were prepared using a modified room-temperature protocol by Tu et al. 2016. 50 mL of LB medium in a 250 mL Erlenmeyer flask was inoculated with 1 mL of the overnight *Pst* culture. Cultivation was carried out in an orbital incubator at 26°C and 220 rpm for 4–6 hours. After reaching an appropriate optical density (OD 0.4–0.6), the cells were centrifuged at 4500 rpm for 10 minutes, the supernatant was discarded, and the cells were gently resuspended in 30 mL of 10% MilliQ glycerol. The washing step was repeated three times. The final resuspension was in 500 μL of 15% MilliQ glycerol, and 100 μL aliquots were stored at –76°C. Electroporation was performed using a Gene Pulser Xcell Electroporation System (Bio-Rad) in 2 mm cuvettes. One aliquot was mixed with plasmid DNA (300–500 ng). Parameters were set to 2100 V, 25 μF, and 200 Ω, with a final pulse duration of about 5 ms. After electroporation, 800 μL LB medium without antibiotics was added. Bacteria were incubated for one hour at 28 °C in an orbital incubator. Finally, transformed *Pst* were plated on LB plates containing kanamycin (50 μg/mL) for selection and incubated for 2 days at 26°C.

### Fumonisin B1 treatment and Trypan Blue staining

Leaves from adult rosettes (No 7–10) were syringe infiltrated with 1 μM FB1 (F1147, Sigma Aldrich) and left for 3 days in growth chambre (21 °C and 10/14 h). Leaves were then cut off, stained by lactophenol Trypan Blue solution by heating for 5 minutes /90 °C, and left for additional 30 minutes to stain. Leaves were subsequently destained in ethanol for 24h, and after that with distilled water. After the last wash, 10% glycerol was added and images of leaves taken. Intensity of trypan blue staining was quantified using Fiji (with normalization toward the background signal; Schindelin et al., 2012). Obtained data were analysed and graphs with statistical evaluation created using GraphPad.

### Microscopy

Imaging was performed at Zeiss LSM880 confocal microscope with 40× water immersion objective, recommended excitation and filters setting for GFP and RFP were used. The images were analysed using Zen 2.1 Software (Carl Zeiss GmbH).

### Text editing

OpenAI’s ChatGPT was used for language editing and refinement in the preparation of this manuscript.

## Acknowledgment

We would like to thank Jana Kaněrová for the technical assistance and Kateřina Malinská for assistance with microscopy.

## Financing

This work was supported by the Ministry of Education, Youth and Sports (MEYS) project TowArds Next GENeration Crops, reg. no. CZ.02.01.01/00/22_008/0004581 of the ERDF Programme Johannes Amos Comenius. We acknowledge the Imaging Facility of the Institute of Experimental Botany AS CR supported by the MEYS CR (LM2023050 Czech-BioImaging), the Czech Academy of Sciences and IEB AS CR.

