## Supplementary tables and figures for "Loss of PR1 function enhances Arabidopsis resistance to *Botrytis cinerea*"

**Supplementary Table 1.** Primers used for creation of *pr1* CRISPR mutant.

| | |
|---|---|
| DT1-BsF-PR1G2 | ATATATGGTCTCGATTGTCCCATGCAGTGGGACGAGGTT |
| DT1-F0-PR1G2 | TGTCCCATGCAGTGGGACGAGGTTTTAGAGCTAGAAATAGC |
| DT2-R0-PR1-G4 | AACACGCCTACCGCTCCTCGTGCAATCTCTTAGTCGACTCTAC |
| DT2-BsR-PR1-G4 | ATTATTGGTCTCGAAACACGCCTACCGCTCCTCGTGCAA |
| PR1ampCRISPR_F | AAGCGATGTTTACGAACCCC |
| PR1ampCRISPR_R | GATTCTCGTAATCTCAGCTCT |
| PR1-gRNA4and2_R | CACAAACTCCATTGCACGTG |
| CAS9 fw | CTCGACTCACGGATGAACACTAA |
| CAS9 rv | AAATCTGCTCAATGATCTCGTCGA |

**Supplementary Table 2.** Primers used for creation of mCherry-PR1 constructs.

| | |
|---|---|
| PP937_pAtPR1-F_EcoRI | ATAGAATTCATATATAACGATCATTGATTAGTATATATAC |
| PP938_PR1-SP-R_NcoI | TCACCATGGCTTTCGAGGGAAGAACAAG |
| PP699_PR1-C-F_ohXFP-NS | GCATGGACGAGCTGTACAAGCAAGATAGCCCACAAGATTA |
| PP939_PR1-3UTR_XhoI | ATACTCGAGGAACATAAGAAATATTGTTTTTGTATTAC |
| PP1166_PR1-C4A-S_forMP | GTTGCAACGCTGCAGCAGCTGGGAATTATGTGAACGAGA |
| PP1165_PR1-Cdel11-S_forMP | CCATAATCAGTTGCAACTATGATTAATGAAGTAATGATGTGATCA |

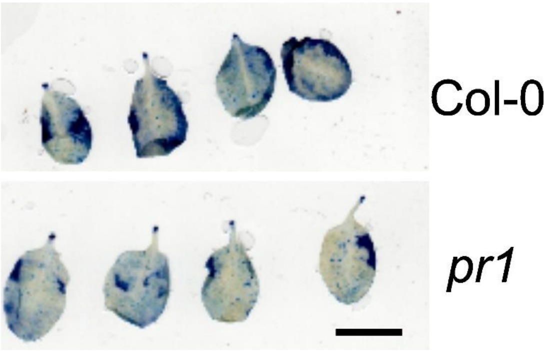


**Supplementary figure 1.** Examples of Trypan Blue-stained leaves after Fumonisin B1-treatement of WT/Col-0 and pr1 mutant plants; bar, 1 cm.

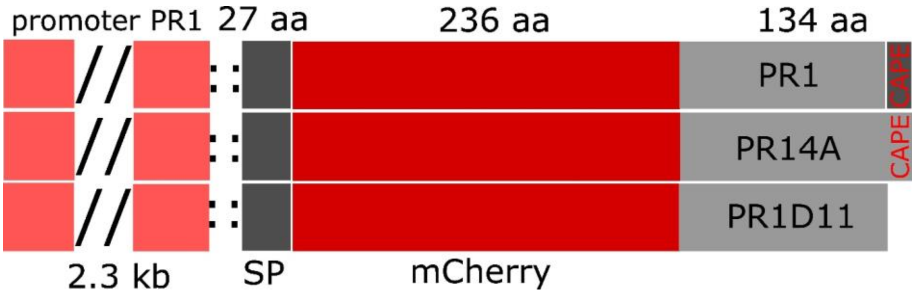


**Supplementary figure 2.** Schematics of designed constructs used for microscopy and complementation assays.

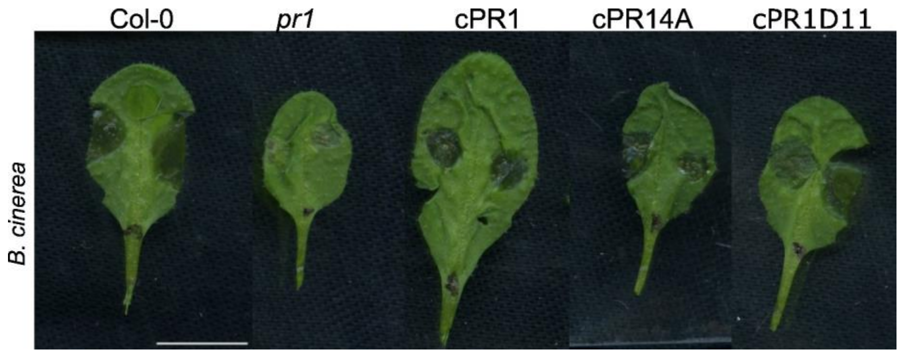


**Supplementary figure 3.** Typical appearance of lesions of *B. cinerea* in Col-0, pr1 mutant and mutant complemented with PR1, PR14A nd PR1D11. Bar, 1 cm.
